# M_1_ selective muscarinic allosteric modulation enhances cognitive flexibility and effective salience in nonhuman primates

**DOI:** 10.1101/2022.10.05.511029

**Authors:** Seyed A. Hassani, Adam Neumann, Jason Russell, Carrie K. Jones, Thilo Womelsdorf

**Author notes:** **Corresponding Author:**; Vanderbilt University, Psychology Department, 301 Wilson Hall, 111 21st Avenue South, 37240-1103 Nashville TN.

## Abstract

Acetylcholine (ACh) in cortical neural circuits mediates how selective attention is sustained in the presence of distractors and how flexible cognition adjusts to changing task demands. The cognitive domains of attention and cognitive flexibility might be differentially supported by the M_1_ muscarinic cholinergic sub-receptor. Understanding how M_1_ mechanisms support these cognitive subdomains is of highest importance for advancing novel drug treatments for conditions with altered attention and reduced cognitive control including Alzheimer’s disease or schizophrenia. Here, we tested this question by assessing how the subtype selective M_1_-receptor specific positive allosteric modulator (M_1_ PAM VU0453595) affects visual search and flexible reward-learning in nonhuman primates. We found that allosteric potentiation of the M_1_ receptor enhanced flexible learning performance by improving extra-dimensional set shifting, by reducing latent inhibition of previously experienced distractors, and by reducing response perseveration in the absence of adverse side effects. These pro-cognitive effects occurred in the absence of apparent changes of attentional performance during visual search. In contrast, non-selective ACh modulation using the acetylcholinesterase inhibitor donepezil improved attention during visual search at doses that did not alter cognitive flexibility and that already triggered gastrointestinal cholinergic side effects. These findings illustrate that M_1_ positive allosteric modulation enhances cognitive flexibility without affecting attentional filtering of distraction, consistent with M_1_ activity boosting the effective salience of relevant over irrelevant objects. These results suggest that M_1_ PAMs are versatile compounds for enhancing cognitive flexibility in disorders spanning schizophrenia and Alzheimer’s diseases.

**Statement of significance:** Muscarinic receptors mediate the pro-cognitive effects of acetylcholine, but it has remained unclear whether they differentially affect the cognitive subfunctions of attentional filtering, set shifting, and learning. To clarify the functional specificity of M_1_ receptors, we assessed these diverse functions using a recently developed, highly selective M_1_ PAM. This M_1_ PAM caused domain-specific cognitive improvement of flexible learning and extra-dimensional set shifting, reduced perseverations and enhanced target recognition during learning without altering attentional filtering functions. These domain-specific improvement contrasted to effects of a non-selective acetylcholinesterase inhibitor that primarily enhanced attention and caused dose limiting adverse side effects. These results demonstrate domain-specific improvements of cognitive flexibility suggesting M_1_ PAMs are versatile compounds for treating cognitive deficits in schizophrenia and Alzheimer’s disease.

## Introduction

Cholinergic activity has far reaching consequences on attention and attentional control functions (1, 2) with long-standing suggestions that cholinergic modulation is involved in faster updating of expectations during learning (3–5). Depleting cholinergic innervation to the prefrontal cortex compromises while stimulation of cholinergic activity can enhance attentional control functions (6–10). These cholinergic effects have been suggested to be supported differently by nicotinic versus muscarinic receptors (11, 12). Antagonizing muscarinic cholinergic action with scopolamine in healthy humans and nonhuman primates (NHPs) increases false alarm rates and impairs sustained attention by slowing response times and impairing signal detection in two-alternative choice tasks (13–17). Consistent with these behavioral effects, neuronal recordings in the prefrontal cortex of NHPs has shown that attentional signaling depends on muscarinic receptor activation (18). One key open question from these insights is to what extent are attentional subcomponent processes supported by muscarinic signaling and whether there are sub-receptors of the muscarinic receptor family that differentially support separable subcomponent processes underlying attention, such as filtering of distracting information, enhancing the flexible updating and shifting of attention sets, or supporting robust goal representations during goal-directed behavior. Each of these diverse subcomponent processes has been associated in prior studies with the muscarinic M_1_ muscarinic receptor, which is widely expressed in the cortex, striatum and hippocampus (19, 20) and may thus mediate some of these muscarinic pro-cognitive effects (2, 21).

One set of prior studies has implicated the M_1_ receptor to memory processes because M_1_ selective drugs can restore deficits in novel object recognition (22, 23), and can partially reverse scopolamine induced deficits in contextual fear conditioning consistent with M_1_ selective compounds enhancing the salience of the (aversive) outcomes during learning (22, 24, 25). But it has remained unclear whether the effects described in these studies are best accounted for by an improvement of memory, or whether enhanced cognitive control processes contribute to more effective encoding of stimuli as opposed to enhancing learning processes. A similar question about the specific cognitive process that are modulated arises when considering the M_1_ effects on different forms of attentional performance. While some studies have shown that M_1_ receptor modulation is important for attentional modulation of neural firing (18, 26), behavioral studies using M_1_ selective ago PAMs (positive allosteric modulators with partial agonistic properties) in NHPs (27) and rodents (28) have not found primary improvements of sustained attention performance. Rather than modulating attention, the M_1_ receptor actions improved behavior only in more demanding task conditions in which M_1_ modulation enhanced the adjustment of performance when task requirements changed (28). These results are consistent with findings showing that M_1_ specific drugs can enhance the likelihood of subjects to apply complex sensorimotor transformations to reach a goal (as in Object Retrieval Detour Tasks) (29), and to facilitate odor-based reversal learning of objects (30). These cognitive enhancing effects suggest that M_1_ receptors may be particularly important for higher cognitive control processes that go beyond attentional focusing or the filtering of distraction (2). However, it is not apparent which particular control processes might be supported by M_1_ receptors as the existing studies used widely varying tasks and a study using one of the classical cognitive control task (the anti-saccade task) was not successful in identifying neural correlates of M_1_ receptor specific effects in the prefrontal cortex of NHPs (31).

The current study set out to address these questions about the specific pro-cognitive role of the M_1_ receptor in supporting cognition. Firstly, to distinguish cognitive subcomponent processes we devised two tasks. A visual search task allowed for distinguishing attentional subcomponent processes by varying distractor load and perceptual interference. And a intra-/extra-dimensional set shifting learning task distinguishing cognitive control processes and cognitive flexibility. Secondly, we assessed NHP performance in these tasks using VU0453595, which is a recently developed subtype selective positive allosteric modulator (PAM) for the M_1_ muscarinic receptor that has exceptional specificity and effectivity to potentiate cholinergic signaling without exerting direct agonistic effects (22, 32, 33). This M_1_ selective PAM is from a family of advanced drug compounds that avoid dose limiting side effects which are prevalent with existing orthosteric compounds (34, 35), and which has the potential to treat deficits in attention control and cognitive rigidity prevalent in schizophrenia, Alzheimer’s disease and substance use disorders (36–40). Assessing the pro-cognitive profile of VU0453595 for these higher cognitive functions is therefore pivotal to advance therapeutic solutions for these widespread neuropsychiatric conditions (22, 41, 42).

We found that the M_1_ PAM VU0453595 exerts an inverted-U shaped improvement of cognitive flexibility, causing faster learning, extra-dimensional set shifting, and reduced perseverations (i.e. enhanced flexibility), while leaving attentional filtering during visual search unaffected. These results are contrasted to the non-selective acetylcholine esterase (AChE) inhibitor donepezil which improved attentional filtering with only marginal effects on cognitive flexibility (43).

## Results

Each of the four monkeys completed 60 sessions composed of 40 vehicle days and 7 days for each of the three tested doses of VU0453595 (0.03, 0.1, and 0.3 mg/kg). No adverse effects were observed in any of the 21 drug dosing days in any of the monkeys.

### M_1_ PAM VU0453595 enhances learning

Animals performed and consistently completed all 21 blocks of the feature learning task per session in all experimental conditions and expectedly showed faster learning in the easier low distractor load condition versus the more difficult high distractor load condition (**Fig. 1C,D**). Administration of VU0453595 improved multiple measures of the learning performance compared to the vehicle control condition. In order to reveal any temporally specific effects on learning, we implemented a linear mixed effects model on the median trials-to-criterion for the feature-reward learning (FRL) task (see **Supplemental Methods**). Faster learning with 0.1 mg/kg dosing was evident in the early, middle, and last thirds of the 21 learning blocks per session with the first third of blocks showing the strongest effects (0.1 mg/kg fixed effect: t(3674) = −2.67, p = .008; first third Cohen’s d = -.228; overall Cohen’s d = -.061) (**Supplementary Fig. S1A**). For this reason, all future analyses of the FRL task use the first third of blocks. Faster learning was particularly evident at low distractor load for which animals reached the trials-to-criterion at 7.93 (SE: 0.81) trials after a block switch with 0.1 mg/kg, compared to 11.03 (SE: 0.38) trials with vehicle (F(3,691) = 3.54, p = .01; η^2^ = .015; Tukey’s, p = .028; Cohen’s d = -.352) (**Fig. 1E**). After the performance criterion was reached, VU0453595 also enhanced plateau performance (**Supplemental Fig. S1B**) and increased the proportion of blocks in which the animals reached the learning criterion at the 0.1 mg/kg dose (**Supplemental Fig. S1C**).

**Figure 1.**
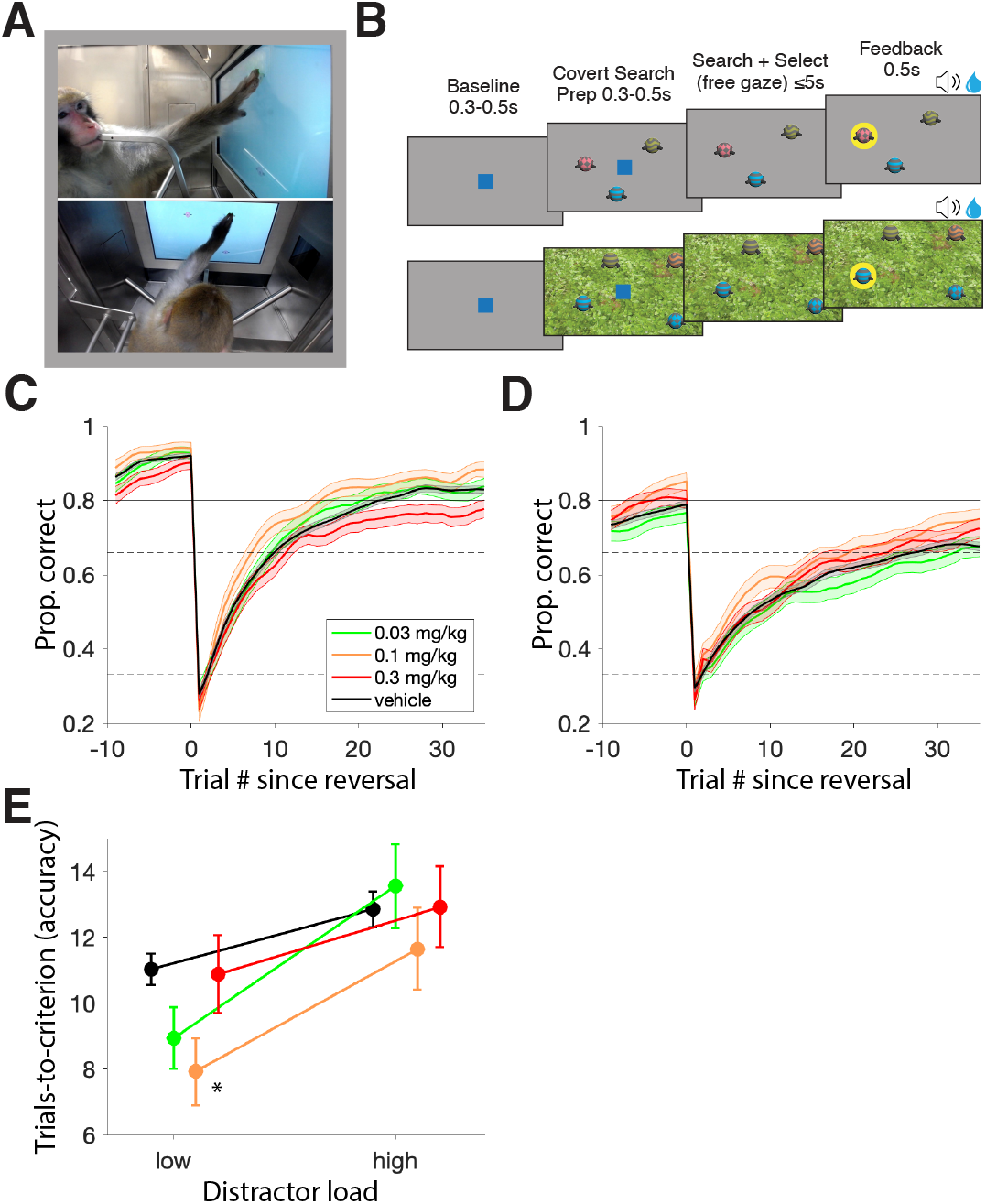
Task design and feature-reward learning task performance enhancement by VU0453595. (**A**) Images of cage-mounted kiosk and monkey Ba utilizing the touch screen to perform the feature-reward learning (FRL) task taken via the video monitoring system. Both images are taken from the same time point from different angles. (**B**) Trial progression of the FRL task (top) and the VS task (bottom). The example FRL task here is a high distractor load block where objects vary in both color and pattern. Although the red checkered object was correctly chosen in this trial, the animal would need to learn through trial-and-error if future red or checkered objects will be rewarded. The example VS block here shows a 3 distractor trial with the target object consisting of the features: blue, striped and straight arms. The red distractor has zero features in common with the target, the yellow distractor has one feature in common with the target (striped pattern) and the blue distractor has two features in common with the target (blue color and straight arms). Trials in either task were initiated by a 0.3-0.5s touch and hold of a central blue square (3° visual radius wide) after which the square disappears (for 0.3-0.5s) and task objects (2.5° visual radius wide) are presented on screen. For the VS task, the background changes to one of 5 images cuing the animal to the task rule while the background remains neutral (gray) in the FRL task. In either task, subjects have 5s to select one of the objects with a 0.2s touch and hold. Failure to choose an object resulted in an aborted trial which was ignored. Feedback for choice selection was provided 0.2s after object selection for 0.5s via both a visual halo around the chosen object as well as a auditory cue alongside any earned fluid. Both the frequency of the audio feedback and color of the feedback halo differed based on outcome. (**C**) Block-wise average learning curves for the low distractor load blocks of the FRL task aligned to block start for vehicle, 0.03, 0.1 and 0.3 mg/kg VU0453595, smoothed after the first 3 trials with a sliding window (shaded area: SE). Dotted horizontal lines signify 0.33 and 0.66 probabilities with the solid horizontal line at 0.8 signifying the block learning criterion. (**D**) The same as C but for the high distractor load blocks. (**E**) Median trials-to-criterion for the low and high distractor load blocks of the FRL task. For the low distractor load blocks, trials to criterion were 11.03 (SE: 0.38), 8.94 (SE: 0.75), 7.93 (SE: 0.81) and 10.88 (SE: 0.94) for vehicle, 0.03, 0.1 and 0.3 mg/kg doses of VU0453595 respectively. Only the 0.1 mg/kg dose was significantly different from vehicle (F(3,691) = 3.54, p = .01; η^2^ = .015; Tukey’s, p = .028; Cohen’s d = -.352). For the high distractor load blocks, trials to criterion were 12.85 (SE: 0.43), 13.56 (SE: 1.03), 11.65 (SE: 1.00), and 12.92 (SE: 0.98) for vehicle, 0.03, 0.1 and 0.3 mg/kg doses of VU0453595 respectively with no significant effect (F(3,565) = .40, n.s.).

Faster learning and improved performance accuracy in the 0.1 mg/kg dose condition was accompanied by faster response times (RTs). Over the course of a learning block, subjects showed a characteristic change of RTs with fast RTs early in the block that slowed down and plateaued around the trial within the block when animals started learning the rewarded target feature (**Fig. 2A,B**). Notably, administering the middle (0.1 mg/kg) dose of VU0453595 led to significantly faster RTs of 870 ms (SE: 23 ms; low load) and 960 ms (SE: 23 ms; high load) relative to vehicle RTs of 960 ms (SE: 11 ms; low load) and 984 ms (SE: 11 ms; high load) (F(3,1672) = 2.97, p = .03; η^2^ = .005; Tukey’s, p = .04; Cohen’s d = -.350) (**Fig. 2C**). Moreover, the number of trials needed for the RTs to plateau was significantly fewer with the 0.1 mg/kg dose taking until trial 6.5 (SE: 0.5) relative to vehicle until trial 8.7 (SE: 0.3) (F(3,193) = 2.67, p < .05; η^2^ = .040; Tukey’s, p = .03; Cohen’s d = -.674) (**Fig. 2D**).

**Figure 2.**
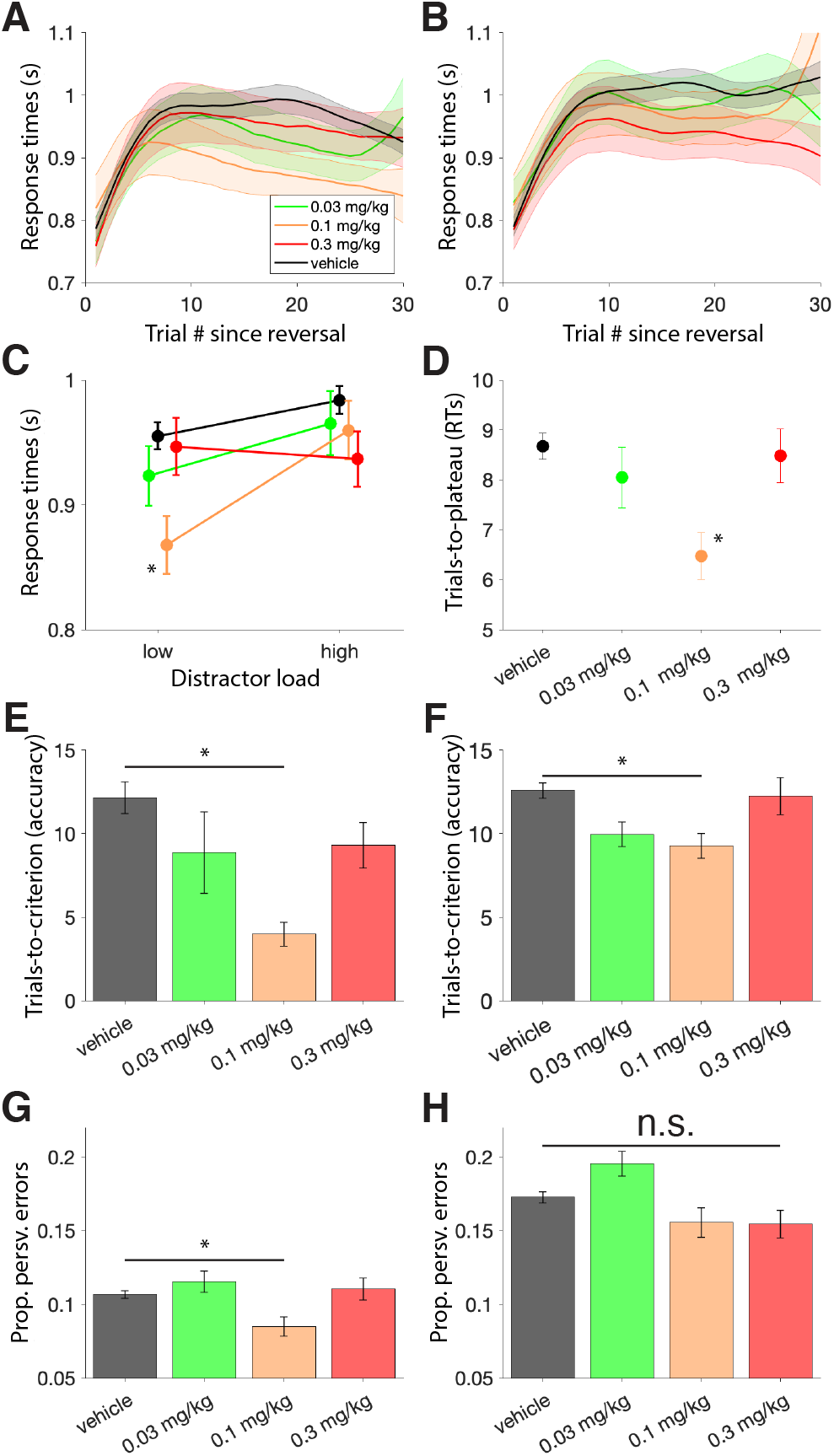
Feature-reward learning task efficiency and cognitive flexibility improvements with VU0453595. (**A**) The average RT curve of each session (correct trials only) aligned to block start for the low distractor load blocks of the FRL task for vehicle, 0.03, 0.1 and 0.3 mg/kg doses of VU0453595 (shaded area: SE) (**B**) The same as A but for high distractor load blocks of the FRL task. (**C**) Block-wise averages of the traces plotted in A and B visualized to compare RTs between distractor load conditions. Low distractor load blocks had RTs of 960 ms (SE: 11), 923 ms (SE: 24), 870 ms (SE: 23) and 974 ms (SE: 23) for vehicle, 0.03, 0.1 and 0.3 mg/kg doses of VU0453595 respectively. High distractor load blocks had RTs of 984 ms (SE: 11), 965 ms (SE: 26), 960 ms (SE: 23) and 937 ms (SE: 22). Only the 0.1 mg/kg dose of VU0453595 was significantly different from vehicle (F(3,1672) = 2.97, p = .03; η^2^ = .005; Tukey’s, p = .04; Cohen’s d = -.350) (**D**) Trials-to-plateau for RTs in the low distractor load blocks defined as the first trial per block (excluding trial 2) that RTs become faster (error bars: SE). Trials-to-plateau was 8.7 (SE: 0.3), 8.0 (SE: 0.6), 6.5 (SE: 0.5) and 8.5 (SE: 0.5) for vehicle, 0.03, 0.1 and 0.3 mg/kg doses of VU0453595 respectively. Only the 0.1 mg/kg dose was significantly different from vehicle (F(3,193) = 2.68, p < .05; η^2^ = .040; Tukey’s, p = .017; Cohen’s d = -.674). (**E**) Block-wise average trials-to-criterion after extra-dimensional shifts were 12.2 (SE:1.0), 8.9 (SE: 2.4), 4.0 (SE: 0.7) and 9.3 (SE: 1.4) for vehicle, 0.03, 0.1, and 0.3 mg/kg doses of VU0453595 respectively. Only the 0.1 mg/kg dose showed a significant difference from vehicle (F(3,122) = 3.15, p = .03; η^2^ = .072; Tukey’s, p = .02; Cohen’s d = -.868). (**F**) Block-wise average trials-to-criterion after intra-dimensional shifts were 12.6 (SE: 0.5), 10.0 (SE: 0.7), 9.3 (SE: 0.7) and 12.3 (SE: 1.1) for vehicle, 0.03, 0.1, and 0.3 mg/kg doses of VU0453595 respectively. Only the 0.1 mg/kg dose showed a significant difference from vehicle (F(3,518) = 3.26, p = .02; η^2^ = .019; Tukey’s, p = .04; Cohen’s d = -.349). (**G**) The proportion of errors that were perseverative in nature with the feature that was perseverated being from the same feature dimension as the target feature. The proportion of perseverative errors from the target feature dimension were 10.7% (SE: 0.2), 11.5% (SE: 0.7), 8.5% (SE: 0.6) and 11.0% (SE: 0.7) for vehicle, 0.03, 0.1, and 0.3 mg/kg doses of VU0453595 respectively with only the 0.1 mg/kg dose being significantly different from vehicle (F(3,1679) = 3.74, p = .01; η^2^ = .007; Tukey’s, p = .01; Cohen’s d = -.243). (**H**) The same as G but with the feature that was perseverated being from the distracting feature dimension (different from the target feature dimension). The proportion of perseverative errors from the distracting feature dimension were 17.3% (SE: 0.3), 19.6% (SE: 0.8), 15.6% (SE: 1.0) and 15.6% (SE: 1.0) for vehicle, 0.03, 0.1, and 0.3 mg/kg doses of VU0453595 respectively. There was a non-significant trend for a main effect of experimental condition (F(3,844) = 2.36, p = .07; η^2^ = .008).

### Improved cognitive control with M_1_ PAM VU0453595

Learning a new feature-reward rule following a block switch entailed either identifying a target feature that was new or from a different feature dimension as in the previous block (extra-dimensional switches, ED), or from the same feature dimension as the previous target (intra-dimensional switches, ID). We found that the 0.1 mg/kg dose with VU0453595 significantly improved learning for both, ED and ID switches (**Fig. 2E,F**) but not switches where the current target was from a novel feature dimension (data not shown). A large improvement was evident for ED switches with the average trials-to-criterion of 4.0 (SE: 0.7) after 0.1 mg/kg dose administration being significantly lower than the average 12.2 (SE: 1.0) trials-to-criterion of the vehicle condition (F(3,122) = 3.15, p = .03; η^2^ = .072; Tukey’s, p = .02; Cohen’s d = -.868) (**Fig. 2E**). Please note that ED switches reported in our task were to a target of the previous distractor feature dimension and thus required disengaging from that dimension in addition to identifying the newly rewarded dimension. ID switches had a more moderate but still significant advantage after administration of 0.1 mg/kg dose of VU0453595 with a trials-to-criterion of 9.3 (SE: 0.7) relative to 12.6 (SE: 0.5) with vehicle (F(3,518) = 3.26, p = .02; η^2^ = .019; Tukey’s, p = .04; Cohen’s d = -.349) (**Fig. 2F**).

A learning advantage after ED and ID switches indicates that VU0453595 at the 0.1 mg/kg dose enhanced cognitive control. Cognitive control processes also entail the ability to avoid erroneous perseverative responding. We quantified the perseverative responses as the proportion of repeated unrewarded choices to a feature in the target-feature dimension or in distractor-feature dimensions. We found that VU0453595 reduced perseverative responding to other features in the target feature dimension at 0.1 mg/kg from the 10.7% (SE: 0.2) of vehicle down to 8.5% (SE: 0.6) (F(3,1679) = 3.74, p = .01; η^2^ = .007; post-hoc analysis of 0.1 mg/kg condition Tukey’s, p = .01; Cohen’s d = -.243) (**Fig. 2G**). Perseverative responding to objects with features of the distractor dimension was moderately, but non-significantly reduced with VU0453595 (F(3,844) = 2.36, p = .07; η^2^ = .008) (**Fig. 2H**).

### M_1_ PAM VU0453595 has no consistent effect on interference control

Cholinergic compounds modulate attention and interference control (12, 43, 44). We evaluated these functions using a visual search (VS) task that varied the requirements to control interference from increasing numbers of distractor objects during search, and from increasing the number of features that were shared between target and distractors (target-distractor similarity, see **Methods**).

Animals showed prominent slowing of target detection times with increasing number of distractors from 3, 6, 9 to 12. VU0453595 did not consistently modulate this slowing with increasing distractor set size (defined as slope of the linear fit) (**Fig. 3A,B**). Similar to target detection times, accuracy was not consistently affected by VU0453595 with no modulation of set size effects. For both the raw values of target detection times and accuracy, some significant changes were observed (see **Supplemental Results**) but no systematic pattern could be extracted (**Fig. 3C,D**). Similarly, VU0453595 did not consistently alter perceptual interference operationalized as changes in performance with increasing similarity between the target and distractors (**Fig. 3E,F**). We did not observe significant changes in the set size effect for target detection times (first block: F(3,236) = 0.54, n.s.; second block: F(3,236) = 1.81, n.s.; **Fig. 3E**) or performance accuracy (first block: F(3,236) = 0.53, n.s.; second block: F(3,236) = 0.51, n.s.; **Fig. 3F**). Similar to the distractor effect, the comparisons of how perceptual interference impacted raw target detection times and performance showed no systematic improvements (see **Supplemental Results**). No changes to speed of processing, operationalized as the time to response during familiarization trials (see **methods**) were observed with VU0453595 at any dose for neither the first VS block (F(3,236) = .56, n.s.) nor the second block (F(3,236) = .35, n.s.) (**Fig. S2**).

**Figure 3.**
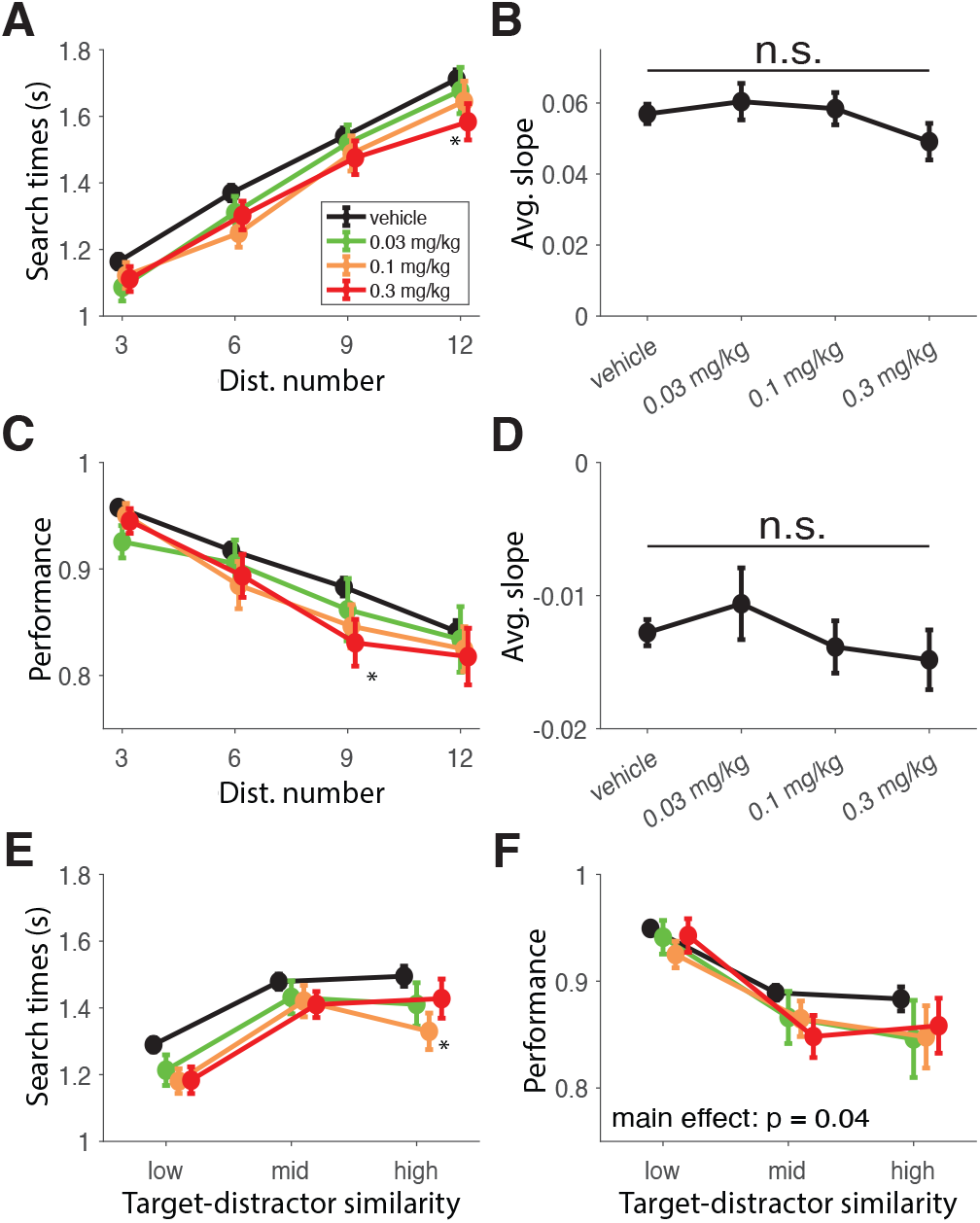
Distractor effect and interference control are not consistently impacted by VU0453595. (**A**) Target detection durations (reaction times) as a function of distractor number for the second VS block. There was a significant main effect of experimental condition with a significant different between the 0.3 mg/kg dose of VU0453595 compared with vehicle (F(3,944) = 3.67, p = .01; η^2^ = .008; Tukey’s, p < .05). The 0.3 mg/kg dose improved search times from 1.16 s (SE: 0.02), 1.37 s (SE: 0.02), 1.54 s (SE: 0.03) and 1.72 (SE: 0.03) with vehicle to 1.11 s (SE: 0.04), 1.30 s (SE: 0.04), 1.48 s (SE: 0.05) and 1.58 s (SE: 0.05) for 3, 6, 9 and 12 distractors respectively. There was no significant change in the first VS block (data not shown). (**B**) The set size effect, operationalized as the slope of the linear fit of search times as a function of distractor numbers for the second VS block (0.057 (SE: 0.003), 0.060 (SE: 0.005), 0.058 (SE: 0.005) and 0.049 (SE: 0.005) for vehicle, 0.03, 0.1 and 0.3 mg/kg doses of VU0453595) was not significant (F(3,236) = .67, n.s.). There was also no significant set size effect in the first VS block (data not shown). (**C**) VS task performance as a function of distractor number for the first VS block. There was a significant main effect of experimental condition with a significant different between the 0.3 mg/kg dose of VU0453595 compared with vehicle (F(3,944) = 3.80, p = .01; η^2^ = .010; Tukey’s, p = .04). The 0.3 mg/kg dose reduced performance from 95.8% (SE: 0.4), 91.8% (SE: 0.7), 88.3% (SE: 0.8) and 84.1% (SE: 1.0) with vehicle to 94.5% (SE: 1.2), 89.4% (SE: 2.0), 83.1% (SE: 2.2) and 81.8% (SE: 2.7) for 3, 6, 9 and 12 distractors respectively. There was no significant change in the second VS block (data not shown). (**D**) The set size effect, operationalized as the slope of the linear fit of performance as a function of distractor numbers for the first VS block (−0.013 (SE: 0.001), −0.011 (SE: 0.003), −0.014 (SE: 0.002) and −0.015 (SE: 0.002) for vehicle, 0.03, 0.1 and 0.3 mg/kg doses of VU0453595) was not significant (F(3,236) = .60, n.s.). There was also no significant set size effect in the second VS block (data not shown). (**E**) VS task search times as a function of target-distractor similarity for the second VS block. There was a significant main effect of experimental condition with a significant different between the 0.1 mg/kg dose of VU0453595 compared with vehicle (F(3,708) = 4.67, p = .003; η^2^ = .018; Tukey’s, p = .02) but no significant set size effect (F(3,236) = 1.81, n.s.). Search times were faster from 1.29 s (SE: 0.02), 1.48 s (SE: 0.02) and 1.49 (SE: 0.03) with vehicle to 1.18 s (SE: 0.04), 1.42 s (SE: 0.05) and 1.33 s (0.05) for low, medium and high average target-distractor similarity respectively. (**F**) VS task performance as a function of target-distractor similarity for the second VS block. There was a significant main effect of experimental condition but no significant post-hoc comparison was found (F(3,708) = 2.84, p = 0.04; η^2^ = 0.011; Tukey’s, n.s.). We also failed to find a significant set size effect (F(3,236) = .53, n.s.).

### Double dissociation of VU0453595 and Donepezil for cognitive flexibility and interference control

VU0453595 improved learning and reduced perseveration, but without reducing interference from distracting objects and features. This pattern of results contrasts to the effects of non-selective acetylcholine esterase inhibitor donepezil for which a prior study using the same tasks as in the current study found that an optimal dose range improved VS performance but without affecting reward learning and perseveration (43). To quantify this difference, we re-analyzed the performance of reward learning and visual search with donepezil in the prior study using the best-dose for VS improvements (0.3 mg/kg) (43). This comparative approach revealed a double dissociation (**Table 1**). VU0453595 enhanced metrics of learning efficiency and cognitive flexibility but not metrics of interference control during VS, while donepezil made the animals more robust against distraction (improved interference control) during visual search but did so without improving feature-reward learning performance. Furthermore, at this dose, donepezil slowed down response times in the FRL task as well as search times in the VS task and even slowed the speed of processing early, partially as a consequence of dose-limiting side effects that accompanied donepezil. In contrast, VU0453595 at 0.1 mg/kg sped up response times in the FRL task without slowing down VS search times or the speed of processing and without any observable side effects (**Table 1**).

**Table 1.** Comparison of performance metrics with the best-doses of VU0453595 and Donepezil. From the FRL task and VS task, we extracted 5 different performance metrics. Learning efficiency and performance entails the number of trials-to-criterion, plateau performance, proportion of learned blocks, response times and trials-to-response-time-plateau. Cognitive control and flexibility entails perseverative error measures and the role of block switches (e.g. ED and ID) on learning efficiency (trials-to-criterion). Speed of processing is a single measure extracted from familiarization trials. Distractor interference entails search time and performance changes as a function of the number of distractors. Perceptual interference entails search time and performance changes as a function of target-distractor similarity.

|  | Extracted Measure | Donepezil<br>(0.3 mg/kg) | VU0453595<br>(0.1 mg/kg) |
| --- | --- | --- | --- |
| Learning Task | Learning efficiency & performance | ↓ | ↑↑↑ |
|  | Cognitive control & flexibility |  | ↑↑↑ |
| Attention Task | Speed of processing | ↓ |  |
|  | Distractor interference | ↑↑↓ | ↑<sup>†</sup> |
|  | Perceptual interference | ↑↓ |  |

†: no systematic effect of 0.1 mg/kg of VU0453595 was found in the VS attention task.

## Discussion

Here, we found that healthy adult rhesus monkeys show M_1_-receptor specific improvements of cognitive flexibility in a feature-reward learning task, while leaving attentional filtering unaffected. In particular, the middle of three doses of the M_1_ PAM VU0453595 increased the speed of learning a new feature-reward rule, particularly with extra-dimensional rule changes. At the same dose animals showed less perseveration on unrewarded features. These pro-cognitive effects contrasted to the absence of distractor dependent changes in accuracy or search times during VS. At the dose range tested no adverse side effects were noted. This result pattern contrasts to the effects of donepezil which improved attentional filtering during VS at a dose at which it did not affect cognitive flexibility, but already resulted in dose-limiting side effects. Taken together, these findings document a functional dissociation of the role of M_1_ receptor modulation with highly selective M_1_ receptor potentiation, suggesting it is a versatile treatment target for disorders suffering from inflexible, rigid cognition and behavior including schizophrenia, Alzheimer’s disease and addiction.

### M_1_ PAM enhances learning and extra-dimensional shifts

We found that the medium dose of VU0453595 improved learning of feature values. Compared to the vehicle control, the medium dose allowed subjects to reach the performance criterion 3.10 trials earlier at the low distractor load condition (**Fig. 1E**) and the number of trials to reach plateau RT decreased by 2.20 trials for low distractor load blocks (**Fig. 2D**) (see **Supplemental Discussion** with regard to dose specificity of the effects). The learning improvement was particularly apparent with extra-dimensional (ED) switches, i.e. when the target feature in a block was from a different feature dimension as the target in the preceding block (**Fig. 2E**). Typically, ED switches take longer and are more difficult than intra-dimensional switches by requiring the recognition of a new dimension and integrating it in a new attentional set (45), suggesting that VU0453595 particularly benefits the flexible updating and switching of attention sets. This finding in NHPs extends the insights that the M_1_ selective ago PAM BQCA (benzylquinolone carboxylic acid) can restore odor-based reversal learning of objects in transgenic mice (30).

Computationally, human and animal studies support the suggestion that the M_1_ PAM might enhance the updating of attention sets. Enhanced learning following ED switches in our task paradigm suggests that the M_1_ PAM allowed the animals to more effectively recognize that previously unrewarded, distracting, features became rewarded. The M_1_ PAM effect is therefore akin to increasing the effective salience of those targets that were ‘learned distractors’ from the previous block while suppressing the salience of current distractor features (**Fig. 1**). Recent modeling suggests that increasing effective salience is achieved with an attention-augmentation mechanism that enhances learning from attended features by actively un-learning (forgetting) unattended features (46). Various studies have documented that such an attention-augmentation mechanism is important for fast learning in complex tasks like the one used here (46–51). The M_1_ PAM effect may thus enhance the effective salience of target features, consistent with neuronal recordings that show M_1_ receptor activation in the prefrontal cortex is necessary during the early processing of targets (52). Support for this suggestion comes from an elegant multi-task study in NHPs that found compromising muscarinic activity with scopolamine increased the pro-active interference of prior spatial information onto current performance in a self-ordered search task (53). The current findings suggest that potentiating M_1_ receptor activity reduces pro-active interference with the net effect of enhanced effective salience.

Recent human studies found that the learning of stimulus-response reward probabilities is enhanced with the AChE inhibitor galantamine (5) and impaired when antagonizing muscarinic receptors with biperiden (54). In a Bayesian framework, these performance improvements were linked specifically to enhanced versus reduced weighting of top-down expectancies and prediction errors during learning (5, 54). In this framework, muscarinic receptor activity determined how fast prediction errors led to belief updating about how stimuli are linked to reward. The results of the current study therefore suggest that enhanced belief updating and effective salience is mediated specifically through the potentiation of the M_1_ receptor. Supporting this conclusion, in rodents, the M_1_ selective ago PAM BQCA reverses scopolamine induced deficits in a contextual fear conditioning consistent with M_1_ enhancing the salience of the (aversive) outcomes during learning (22, 24, 25).

These functions of the M_1_ receptor may be realized in the prefrontal cortex. In primates, reversal learning and the extra-dimensional updating of attentional sets depend on dissociable subareas of the prefrontal cortex with ED shifting and the recognition of attention sets depending particularly on the ventrolateral prefrontal cortex (55, 56). Support for such a prefrontal mechanisms comes from a rodent study that found the M_1_ selective PAM TAK-701 can partially reverse a deficit of target detection selectively on signal trials that followed no-signal trials when the deficit was induced by partially (∼60%) depleting ACh afferents to the prefrontal cortex (28). These considerations support the notion that M_1_ receptors in prefrontal cortex are pivotal for the improved updating of attentional sets (2).

In previous work, faster learners of feature-reward associations were shown to have improved working memory capacity (46), which raises the possibility that M_1_ allosteric modulation may have affected learning through enhanced short-term memory of targets. We believe this is unlikely. While the non-selective muscarinic antagonist scopolamine impairs short term memory retention and non-selective AChE inhibitors partially reverse the deficit (57–61), the short-term deficits can be independent of the delay and more prominent for short or intermediate delays, making it unlikely that muscarinic receptors have primary effects on recurrent persistent delay representations (53, 59, 62, 63).

### M_1_ PAM reduces perseverative responding

A second main result of the current study is VU0453595 reduced response perseveration, allowing animals to avoid repeating erroneous responses to objects with the same non-rewarded features (**Fig. 2E**). This finding supports early insights into the effects of the muscarinic antagonist scopolamine in the prefrontal cortex to increase omissions (64), suggesting that it is the M_1_ muscarinic receptor that is particularly important for minimizing error rates. Support for the M_1_ specificity of these effects also comes from a study treating transgenic mice with an M_1_ selective ago PAM which resulted in reduced erroneous choices of compound object discrimination in the trials after reversing object-reward associations (30).

Perseverative responding is the key characteristic of inflexible, habitual responding because it reflects that performance feedback is not utilized for adjusting behavior. It has been shown that performance feedback triggers transient activation of cholinergic neurons in the basal forebrain in mice (65) and activates the basal forebrain in humans (66). In the prefrontal cortex, cholinergic transients trigger gamma activity (67) that depends specifically on local M_1_ receptors (52).

Taken together, the reduction of perseverative responding with VU0453595 implicates the M_1_ receptors also in the effective processing of feedback to adjust future performance. Perseverative, habitual responding is a hallmark of multiple psychiatric disorders including schizophrenia, obsessive compulsive disorder and substance use disorders (68, 69). The current result therefore bears particular relevance by suggesting that potentiating the M_1_ receptor critically reduces perseverative response tendencies (70).

### M_1_ PAM has no consistent effect on interference control over distractors

We found that VU0453595 did not affect VS performance differently with few or many distractors. Target detection response times were moderately faster and accuracy was moderately lower to a similar extent for 3, 6, 9, or 12 distractors (**Fig. 3A-D**). This finding shows that the M_1_ PAM dose range that improved cognitive flexibility did not alter attentional filtering of distracting information. This finding adds clarity to diverse results in previous studies. Firstly, the absence of M_1_ specific distractor effects resonates with a recent finding in rodents that the M_1_ selective PAM, TAK-071, did not modulate the distracting effects of light on/off switches during a sustained attention task, but started to improve performance in the second half of testing when distraction ended and the animals adjusted to a no-distractor regime (28). This result pattern is congruent with our result pattern. Allosteric modulation of the M_1_ receptor improved adjusting behavior to challenges, but without improving interference control from distraction. A similar lack of effects of muscarinic modulation on distractor interference control were found in other task contexts. Scopolamine-induced deficits of continuous recognition performance can be partially reversed with an M_1_ selective agonist (71) or the non-selective muscarinic agonist milameline (72, 73), but this deficit reversal is independent of the similarity between distracting and target objects (27). Similarly, scopolamine does not alter distractor effects in an attentional flanker task, but rather causes an overall slowing and selective impairment of learning reminiscent of the reward learning effect we found (74).

The observed result pattern with the M_1_ PAM contrasts to apparent effects to reduce distraction with nicotinic modulation (12, 44, 75, 76), with non-selective cholinergic increases using donepezil (43) (see **Table 1**), or with the improvement of target detection accuracy and visuo-spatial attentional orienting when enhancing cholinergic transmission from the basal forebrain (1, 77–79). Particularly relevant in this context is a prior NHP study that found the nicotinic alpha-4/beta-2 receptor agonist selectively enhanced distractor filtering when two stimuli underwent salient changes but had no effect on reversal learning speed (44).

One caveat when interpreting the absence of a drug effect is that we cannot know whether higher VU0453595 doses than were used here would have affected distractor filtering during VS performance. The highest dose used in this study (0.3mg/kg, oral) is a magnitude lower than the ≥3mg/kg doses that previous studies found to be safe and void of adverse side effects (80), suggesting that future studies will need to identify possible dose specific effects on attention functions.

## Limitations

While our study already tested multiple markers of cognitive flexibility and attention, it was not yet incorporating tests of other domains that M_1_-modulating drugs might affect and which are compromised in psychiatric patient populations such as long-term memory and motivation (13, 81–83). Further tasks, where we can extract measures of longer-term memory processes and motivation for example would be important additions for a more comprehensive characterization of possible M_1_-dependent behaviors (see **Supplemental Discussion**). Such an expansion of extracted measures would align well with efforts to develop multi-task batteries for NHPs covering a wide range of cognitive domains (53, 84–87).

## Conclusion

In summary, the M_1_ positive allosteric modulator VU0453595 produced selective improvements in cognitive flexibility in the absence of adverse side effects. The results were obtained with cognitive tasks that tap into real-world cognitive demands for adjusting to the changing relevance of visual objects. This result pattern suggests that M_1_ PAMs will be powerful targets for drug discovery efforts to augment cognitive flexibility.

## Methods and Materials

### Subjects

Four adult male rhesus macaques (*Macaca mulatta*) were separately given access to a cage-mounted Kiosk Station attached to their housing unit where they performed a visual search attention task and a feature-reward learning task via a touchscreen interface (85) (**Fig. 1A**) (see supplemental materials for more details).

### Compounds and Procedures

VU0453595 was synthesized in house (30, 33) and mixed with a vehicle of 18g of strawberry yogurt and 2g of honey provided to the monkeys in a small paper cup (oral administration). All monkeys received vehicle or drug 2h prior to the start of behavioral performance and were observed to ensure full consumption of vehicle or drug. VU0453595 was administered once per week to allow appropriate washout. Based on the weight of each animal, drug volume was calculated for 0.03, 0.1 and 0.3 mg/kg doses. Drug side effects were assessed 15 min following drug administration and after completion of the behavioral performance with a modified Irwin Scale for rating autonomic nervous system functioning (salivation, etc.) and somato-motor system functioning (posture, unrest, etc.) (43, 88–90). Furthermore, monkeys’ behavioral status was video-monitored throughout task performance.

### Behavioral Paradigms

Monkeys performed a sequence of two tasks in a single behavioral session including a VS task block, 21 reward learning task blocks and finally, another visual task block. Rewarded and unrewarded objects in the VS task and FRL task were multidimensional, 3D rendered Quaddle objects (91) that shared few or many features of different features dimensions (colors, shapes, arms, body patterns). The VS task varied the perceptual target-distractor similarity by changing the average number of common features between distractors and the target object. The FRL task varied the complexity of the feature space by varying features of objects in only one or of two feature dimensions from trial to trial.

Animals first performed a VS task block consisting of ten familiarization trials that showed the same object on a screen without distracting objects, followed by a set of 100 trials that contained the previously shown object amongst 3, 6, 9, or 12 distracting objects (**Fig. 1B**). Animals received fluid reward for touching the previously shown target object. Following the VS task, the animals performed a FRL task that required learning, by trial-and-error, which feature was associated with reward for each block of 35-60 trials. Trials in this task always contained 3 objects that each contained one or two features, depending on the block, with only one instance of each feature presented per trial (**Fig. 1B**).

The FRL task indexes cognitive flexibility by testing how fast subjects learn which feature is rewarded when the feature-reward rule switched between blocks. Block switches were un-cued and could involve switching the newly rewarded feature to the same or different feature dimensions, which makes the task similar to conceptual set shifting tasks, but different by using a larger set of features that varied within and across sessions in order to vary task difficulty. In each trial three objects were shown that varied either in features of one feature dimension (e.g. having different colors *or* body shapes), or that varied in features of two feature dimensions (e.g. having different colors *and* body shapes). Choosing the object with the correct feature was rewarded with a probability of 0.85. Blocks where only 1 feature dimension varied (low distractor load) were easier as there were less distracting features than in blocks with 2 varying feature dimensions (high distractor load).

## Statistical Analysis

Data were analyzed with standard nonparametric and parametric tests with test statistics, p values and effect sizes reported where appropriate in text. For detailed statistical methods, please see the **Supplemental Information**.

## Acknowledgements

This work was supported by the National Institute of Mental Health of the National Institutes of Health under Award Number R01MH129641 (TW). The content is solely the responsibility of the authors and does not necessarily represent the official views of the National Institutes of Health.

## Author Contributions

S.A.H., J.R., C.J. and T.W. conceived the experiments. A.N. and S.A.H. performed the experiment. J.R. and C.J. contributed the drug compounds. S.A.H analyzed and visualized the data. S.A.H. and T.W. wrote the first paper draft. All authors contributed writing the paper.

## Supplemental Information

### Supplemental Materials and Methods

#### Subjects

All animal related experimental procedures were in accordance with the National Institutes of Health Guide for the Care and Use of Laboratory Animals, the Society for Neuroscience Guidelines and Policies, and approved by Vanderbilt University Institutional Animal Care and Use Committee.

Four pair-housed adult male rhesus macaques (*Macaca mulatta*), 7-11 years old and weighing ∼8-15 kg were subjects in this experiment. Monkeys in each pair were separately given access to a cage-mounted Kiosk Station attached to their housing unit uniformly at either 11am (monkeys Ig and Ba) or at 1pm (monkeys Re and Si). Each monkey was overtrained and engaged with and completed a visual search attention task and a flexible feature-reward learning task via a touchscreen interface (1) with the software being controlled by the Unified Suite for Experiments (USE) (2).

Of the four monkeys, two (monkeys Si and Ig) had previously been involved in a similar study utilizing the acetylcholine esterase inhibitor donepezil (3) with over 6 months between experiments for washout. Prior to the donepezil experiments, monkey Ig had also previously been exposed to a different experimental M_1_ PAM. Monkeys Ba and Re were naïve to VU0453595 and other neurological or psychiatric medications.

#### Comparison table

The comparison table (Table 1) between the best dose of donepezil and VU0453595 was based on the data collected in a previous study (3) but for all measures, identical methods were applied to both datasets consistent to what is described here.

#### Statistics

Within the feature-reward learning (FRL) task, we use trials-to-criterion to quantify learning efficiency with the criterion being defined as the first trial after at least 1 error which preceded a string of 10 trials with 70% or greater performance. The 70% performance threshold is different from our previous work (3) which was set to 80% performance, which was and still is the criterion for block switches in the FLR task. We found this new threshold to better reflect the occurrence of learning and led to only a mean 0.37 (0 median) trial difference in baseline trials-to-criterion overall. The comparisons between VU0453595 and donepezil were made using the same definition for each measure.

Block switches in the FRL task were labelled based on the status of the target feature relative to the previous block as extra-dimensional, intra-dimensional or as involving a novel target feature dimension. For novel target blocks, the rewarded feature dimension was not present in the previous block, independent of the present or previous block’s dimensionality. Similarly, intra-dimensional shift blocks involved the same rewarded feature dimension but a different rewarded feature (e.g. a different color) as the previous block, independent of their dimensionality. However, for extra-dimensional shift blocks, the previous block must have been a high load (objects varying in 2 feature dimensions) block where the current block’s target feature was from the previous one’s distracting feature dimension. Extra-dimensional shift blocks themselves could be either low or high distractor load.

In the FRL task, for each session, reaction times (RTs) were averaged and smoothed using a 5 trial shifting window for low and high distractor load blocks separately (**Fig. 2A,B**). We then defined the time to plateau as the first trial per session, excluding trial two, where the RTs began to decrease.

Perseverative errors were quantified based on the features of the erroneously chosen object. The consecutive errors could be made with objects containing the same feature from the distracting or target feature dimensions. The proportion of perseverative errors are reported as a percentage of all errors (**Fig. 2G,H**).

In order to account for temporally specific effects on learning efficiency with VU0453595, as seen with other cholinergic compounds (3), we applied a linear mixed effects model (LMEM) to the trials-to-criterion. The LMEM had three main effects: experimental condition, distractor load and temporal bin (thirds), while individual monkeys were treated as random effects:

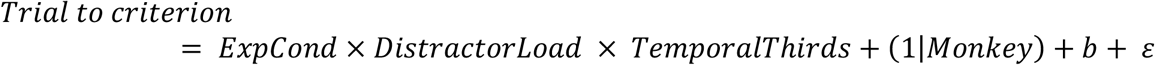

Given the results of the LMEM and the maximal effect size with the first third of blocks in the FRL (**Fig. S1A**), all analyses for the FRL task used only the first third of blocks to capture the period where VU0453595 had its strongest effect on performance.

Effect sizes were reported as either eta squared values when referring to ANOVA results or Cohen’s d when appropriate (i.e. when post-hoc analysis showed a significant effect at a single dose). The Cohen’s d was computed by directly comparing vehicle to the significant dose using this formula:

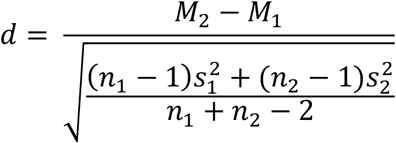

### Supplemental Results

#### Feature-reward learning task

After reaching performance criterion, VU0453595 also resulted in higher plateau accuracy (drug condition main effect: (F(3,1672) = 3.22, p = .02; η^2^ = .005); low distractor load accuracy: 90.9% (SE: 1.9%); 95.9% (SE: 0.9%); 90.9% (SE: 1.8%); 91.0% (SE: 0.7%) for 0.03. 0.1, 0.3 mg/kg and vehicle respectively; high distractor load accuracy: 74.4% (SE: 3.0%); 80.5% (SE: 2.9%); 84.8% (SE: 2.3%); 77.9% (SE: 1.1%) for 0.03. 0.1, 0.3 mg/kg and vehicle respectively) (**Supplemental Fig. S1B**). The middle dose of VU0453595 (0.1 mg/kg) also increased the proportion of blocks in which animals reached the learning criterion (F(3,1369) = 2.93, p = .03; η^2^ = .006; Tukey’s, p = .02). Animals reached the learning criterion of 80% accuracy over 10 successive trials in 90.5% (SE: 2.4; low load) and 72.1% (SE: 3.7%; high load) of blocks in the vehicle condition. Tukey’s HSD multiple comparisons test among proportions revealed that at the 0.1 mg/kg dose, VU0453595 significantly increased the proportion of learned blocks in the low load condition to 98.7% (SE: 2.6%) (p = .04) (**Fig. S1C**).

#### Visual search task

In the first VS block, target detection times across distractor conditions were not different with VU0453595 relative to vehicle control (F(3,944) = 1.67, n.s.; η^2^ = .004), with the exception of faster target detection times in the second VS block at the 0.3 mg/kg dose (experimental condition main effect: F(3,944) = 3.67, p = .01; η^2^ = .008; Tukey’s, p < .05). With regards to performance, in the VS block at the end of the session there were no significant effects, while in the first VS block there was a significant main effect of drug condition (F(3,944) = 3.80, p = .01; η^2^ = .010) with reduced accuracy at 0.3 mg/kg dose (Tukey’s, p = .04) irrespective of the number of distractors.

Despite the lack of set size effects, the raw target detection times were overall significantly faster with the 0.1 mg/kg dose in the second block (F(3,708) = 4.67, p = .003; η^2^ = .018; Tukey’s, p = .02) with more improvement with high target-distractor similarity (cohen’s d = -.447) than low target-distractor similarity (cohen’s d = -.427) (**Fig. 3E**). There was also a general reduction in performance during the first block (F(3,708) = 2.84, p = 0.04; η^2^ = 0.011) (**Fig. 3F**).

### Supplemental Discussion

#### Possible M_1_ agonism

Although *in vitro* data suggests little to no agonistic properties of VU0453595 (4), we cannot completely rule out this possibility in the current study. At high doses, M_1_ PAMs may activate M_1_ receptors, even with little (sub-threshold) or no endogenous ACh. Especially in cells with high M_1_ receptor expression, our macaques may be subject to M_1_ agonism with the highest dose of VU0453595 tested. This may explain why the pro-cognitive effects seen at the middle dose of the three tested doses of VU0453595 are not observed at the highest tested dose. This would suggest that the endogenous signaling at the M_1_ muscarinic receptor supporting cognitive flexibility is sensitive to exogenous intervention which would predict lower efficacy with orthostatic agonists. The possible agonism of M_1_ PAMs such as VU0453595 *in vivo* should be the subject of future studies.

#### Possible contributions of M_1_ potentiation of memory or motivation/effort control

The current study dissociated the relative importance of an M_1_ PAM for cognitive flexibility and attentional filtering and contrasted these effects to those of donepezil (**Table 1**). The functional dissociation of the drug effects highlights the importance of a multi-task paradigm for understanding drug actions on behavior (3, 5, 6) and supports efforts to develop multi-task batteries covering a wide range of cognitive domains (1, 5–8). While our study tested already multiple markers of cognitive flexibility and attention, it was not yet incorporating tests of domains that M_1_ modulating drugs might also affect. For example, scopolamine challenges have long suggested that M_1_ receptors in the medial temporal lobe support longer-term memory processes (9–11), making it possible that M_1_ receptor modulation might have positive consequences in this domain.

Motivation and the ability to control effort are other domains that we did not test and which some studies have suggested to be modulated my mAChRs. The task we deployed varied cognitive load which inevitably increases difficulty and the amount of effort subjects needed to exert. Although we did not control for motivational factors explicitly, visual inspection suggested it was not modulated by VU0453595 because the learning improvements were somewhat more pronounced at lower than higher load in the learning task and did not vary with increasing distractor difficulty (target-distractor similarity) in the search task. These findings resonate with the results of a scopolamine challenge study in NHP that found no effects of increasing difficulty in a stimulus-location association learning task (12). However, when testing for a memory load effect with a visuo-spatial paired associate task, Taffe and colleagues (13) found that scopolamine reduced performance particularly when 3 or 4 stimulus-object associations needed to be learned and retrieved but not when 1 or 2 associations were involved. Such a memory load differs from the cognitive load that we imposed by increasing the number of distracting features in the learning task and from the perceptual load that we varied with increasing target distractor similarity. However, it will be important to identify in future studies which motivation or load dependent processes are modulated specifically by M_1_ selective mAChR modulation.

### Supplemental Figures

**Figure S1.**
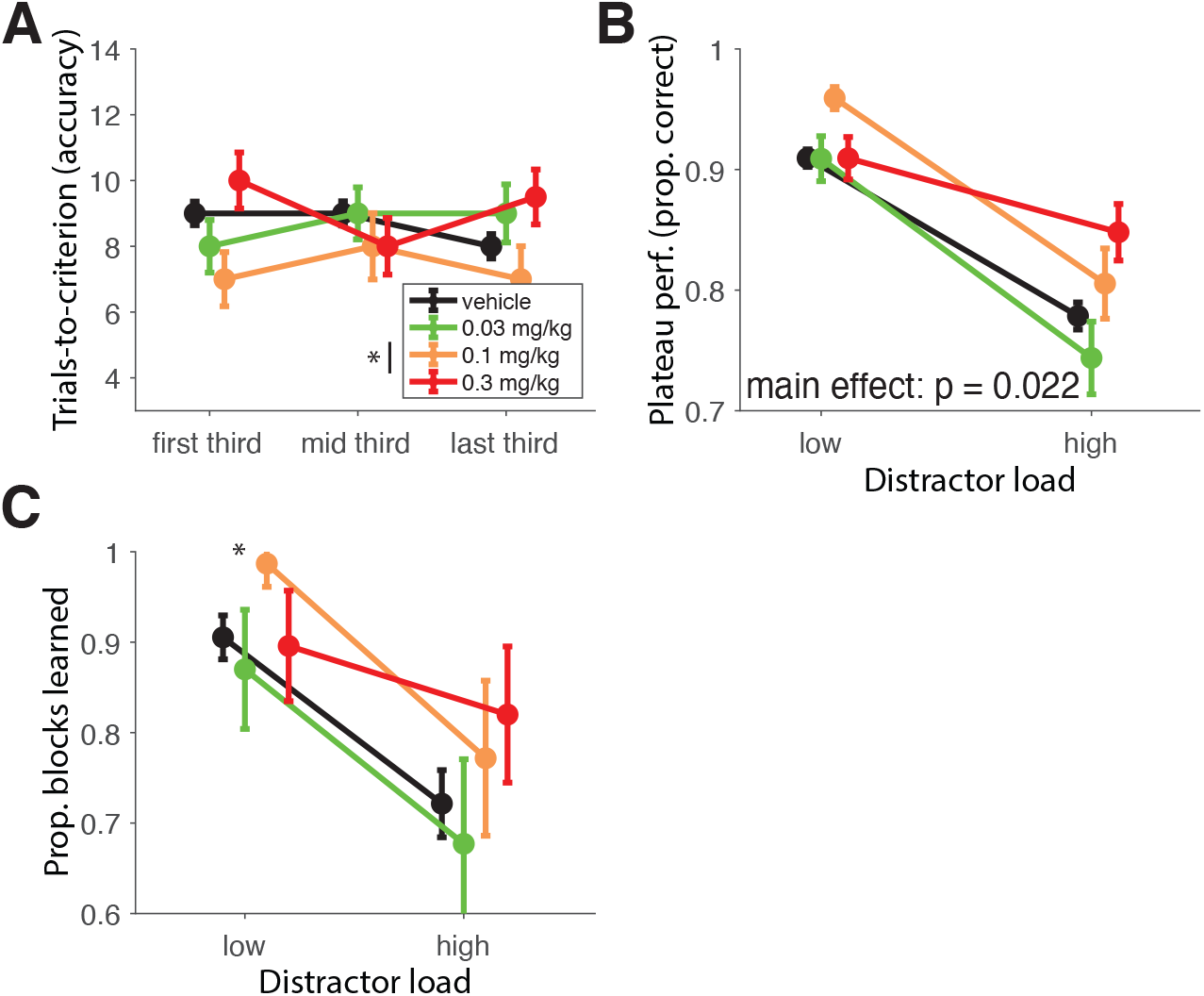
VU0453595 enhances multiple measures of learning performance. (**A**) The median trial-to-criterion, visually combined for low and high distractor load conditions temporally split by their presentation within a session (7 blocks in each third) for vehicle, 0.03, 0.1 and 0.3 mg/kg doses of VU0453595. The LMEM used the experimental condition, temporal bin (thirds) and distractor load as fixed effects. There was significantly faster learning with 0.1 mg/kg which showed the strongest effect size during the first third of FRL blocks (0.1 mg/kg fixed effect: t(3674) = −2.67, p = .008; first third Cohen’s d = -.228; overall Cohen’s d = -.061). (**B**) Average performance in the final 10 trials of low and high distractor load blocks of the FRL task. For the low distractor load blocks, plateau performance was 90.95% (SE: 0.73), 90.90% (SE: 1.86), 95.92% (SE: 0.92) and 90.94% (SE: 1.76) for vehicle, 0.03, 0.1 and 0.3 mg/kg doses of VU0453595 respectively. For the high distractor load blocks, plateau performance was 77.86% (SE: 1.12), 74.37% (SE: 3.01), 80.54% (SE: 2.91) and 84.80% (SE: 2.34) for vehicle, 0.03, 0.1 and 0.3 mg/kg doses of VU0453595 respectively. There was a significant main effect of experimental condition (F(3,1672) = 3.22, p = .022; η^2^ = .005) but post hoc analysis (Tukey’s) showed no single dose as significantly different from vehicle. (**C**) Average proportion of learned blocks (defined as blocks that reached the 70% performance over 10 trials) per session in the FRL task. For the low distractor load blocks, the proportion of blocks learned was 90.54% (SE: 2.42), 87.00% (SE: 6.59), 98.68% (SE: 2.56) and 89.58% (SE: 6.11) for vehicle, 0.03, 0.1 and 0.3 mg/kg doses of VU0453595 respectively. For the high distractor load blocks, the proportion of blocks learned was 72.14% (SE: 3.71), 67.71% (SE: 9.35), 77.17% (SE: 8.58) and 82.00% (SE: 7.53) for vehicle, 0.03, 0.1 and 0.3 mg/kg doses of VU0453595 respectively. Pair-wise comparisons between the VU0453595 doses and vehicle revealed a significant improvement at the low distractor load with the 0.1 mg/kg dose (Tukey’s multiple comparison test among proportions: q = 4.082, q_crit_ = 3.633) and no significant changes at the high distractor load (Tukey’s multiple comparison test among proportions, n.s.).

**Figure S2.**
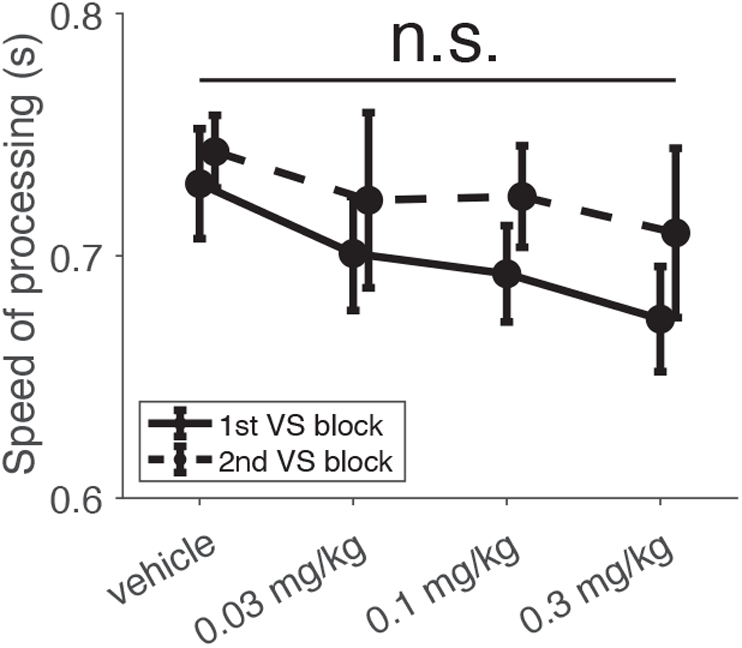
VU0453595 does not impact the speed of processing. The speed of processing for the first and second VS blocks, defined as the time animals took to touch the only object on screen (during familiarization trials). In the first VS block, speed of processing was 0.730 s (SE: 0.023), 0.701 s (SE: 0.024), 0.693 s (SE: 0.020) and 0.674 s (SE: 0.022) for vehicle, 0.03, 0.1 and 0.3 mg/kg doses of VU0453595 respectively. In the second VS block, speed of processing was 0.743 s (SE: 0.015), 0.723 s (SE: 0.036), 0.725 s (SE: 0.021) and 0.710 s (SE: 0.035) for vehicle, 0.03, 0.1 and 0.3 mg/kg doses of VU0453595 respectively. No significant changes were observed for the first (F(3,236) = .56, n.s.) or second VS blocks (F(3,236) = .35, n.s.).

